# Using low-cost alcohol sensors to monitor real-time olfactory information during odor-guided navigation

**DOI:** 10.1101/665703

**Authors:** Mohammad F. Tariq, Suzanne M. Lewis, Aliena Lowell, David J. Perkel, David H. Gire

**Affiliations:** Graduate Program in Neuroscience, University of Washington; Department of Psychology, University of Washington; Departments of Biology & Otolaryngology, University of Washington; University of Washington Institute for Neuroengineering

## Abstract

Olfaction — an evolutionarily conserved chemosensory modality — guides navigation and decision-making in organisms from multiple phyla within the animal kingdom. However, using olfactory cues to guide navigation is a complicated problem. This is because the spread of odor molecules from the source is governed by turbulent and chaotic air currents, resulting in intermittent and spatiotemporally varying sensory cues as odor plumes. Precisely correlating olfactory information with behavior and neurophysiology from freely behaving animals has thus been a challenging avenue due to the dynamic nature of the odor plumes. Current technologies for chemical quantification are cumbersome and not feasible for monitoring the olfactory information at the temporal and spatial scales relevant for plume and odor tracking in animals. Here we present an alternate method for real-time monitoring of olfactory information using low-cost, lightweight sensors that robustly detect common solvent molecules, like alcohols. Paired recordings were made from these ethanol sensors with a Photoionization detector (PID) to precisely controlled ethanol stimuli. The ethanol sensor recordings were then deconvolved using a double exponential kernel, showing robust correlations with the PID recordings at behaviorally relevant time, frequency and spatial scales. Furthermore, the light weight of these sensors allows us to mount them on the heads of freely behaving rodents engaged in odor-guided navigation. Our preliminary experiments in mice show robust behavioral and neurophysiological responses correlated with ethanol plume contacts detected by these sensors.

## Introduction

Odor molecules emanating from a source spread due to turbulent air currents[1,2]. Currents of varying time scales and of different sizes exist in all fluids. Diffusive forces, resulting from Brownian motion, are the primary factors in creating a chemogradient within a layer of static fluid contact with the source. However, this diffusive layer is very narrow, limiting the spread of molecules to very short distances (<1 mm). Furthermore, spread via diffusion takes immensely long times to reach further distances. Above this very narrow diffusive layer, a velocity gradient exists within several layers. The size of currents, and their velocity profile within these layers depend on the characteristics of the bulk flow and atmospheric conditions. These dynamically varying currents result in a turbulent mixing of the odor molecules with the fluid molecules. The net result of these forces is a dynamically varying spread of odor molecules in time and space as odor plumes. Hence, time-averaged concentrations of the odorant in space are a poor measure of the odor profile that a non-stationary searcher, using odors to locate the source will experience[3]. An approach to overcome this problem is to use sensors that monitor the real-time concentration of the odorant molecules as the searcher navigates during plume tracking[4].

Current technologies for chemical quantification are not feasible for mobile operations. Photoionization detection (PID) is a technology that is routinely used by olfactory researchers[5], but the sensor in this case is also relatively big and expensive. A technology that is used by the insect community is Electroantennography (EAG) [5]. However, EAG signals often degrade over the time scale of hours. We thus developed a method using metal oxide gas sensors that are routinely used in environmental monitoring systems as a feasible alternative for mobile monitoring of the environment. One limitation of using the metal oxide sensors has been the long decay time of the sensor recordings in response to transient activation with the chemical stimulus. Here we first conducted paired alcohol sensor and PID recordings to behaviorally relevant stimuli and developed a method to deconvolve the sensor recordings to match the PID recordings. Furthermore, we also present preliminary results correlating the deconvolved signals resulting from the alcohol sensors to neurophysiological responses in head-fixed, and behavioral responses in freely navigating mice. This method can serve as a cost-effective way to correlate real-time olfactory information with behavior and physiological responses and would thus greatly benefit olfactory research.

## Materials and Methods

### Paired PID and ethanol recordings

Figaro TGS 2620 Organic Solvent Vapor Sensor (powered by a 5V DC voltage from an Arduino) was used to monitor the relative concentration of the ethanol in air. To prevent extended contact of odor-laced air with the sensor, the head cap of the sensor was removed. Paired PID and alcohol sensor recordings were then conducted ~10-15 mm downstream of an odor port pushing ethanol vapors from a tube controlled by valve. The PID and sensor recordings were digitized by NI USB-6009 DAQ (National Instruments) at a sampling frequency of 500 Hz. The data acquisition and control of valve was carried out through LabView by custom-written scripts. For recordings in a dynamic plume, paired recordings were carried out downwind of an ethanol port in a custom-designed arena at multiple locations. Data from locations near the port, middle of the arena, and farthest downwind were pooled.

### Deconvolution of the ethanol sensor

The raw alcohol sensor recordings were low-pass filtered using a digital filter in MATLAB (cutoff frequency: 40 Hz). The frequency content in the filtered signal was then obtained by the fft routine. A kernel designed using a difference of two exponentials was then designed (see Results) using the equation:

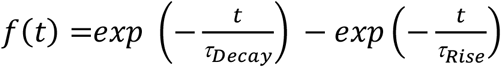

The frequency content of the kernel, obtained by the fft routine, was then used to divide the filtered signal. The resulting spectrum was then converted back into the time domain by taking the inverse fft of the signal. The difference between the PID recordings and the deconvolved signals was calculated by normalizing both the PID and deconvolved signal. This allowed for the comparison of the waveform of the two signals with no particular emphasis on the amplitude of the responses.

## Results

### Difference of two exponentials kernel to deconvolve the raw alcohol sensor responses

To test the feasibility of the ethanol sensor to record behaviorally relevant odor stimuli, we conducted paired PID and ethanol sensor recordings in response to brief ethanol stimuli (**Fig. 1A**). As can be seen, the ethanol sensor recordings take a long time to decay compared to the PID responses. To improve our understanding of the rise time and time of decay of the ethanol sensor responses, the individual recordings were averaged to determine the mean time rise and tau decay. The mean rise and decay signal (**Fig. 1B**) were then exponentially fit to obtain the tau rise (0.06 s) and tau decay (3.5 s). Modeling the ethanol response as a difference of two exponentials [7], we deconvolved individual ethanol sensor recordings using a family of kernels with a range of tau rise and tau decay values. These deconvolved signals were then compared with the PID recordings, taken as the ground truth. The dependence of the mean error between the deconvolved sensor recordings and the PID recordings as a function of tau rise and tau decay values of the kernel is presented in **Fig. 1C**. The optimal tau rise values for the kernel were found to be less than 0.02 s and the tau decay values to be less than 0.5 s. Hence, we set the value of tau rise for the kernel to be 0.002 s and that of tau decay to be 0.5 s. Using this kernel, the raw ethanol recordings presented in **Fig. 1A** were deconvolved and are shown in **Fig. 1D**. For comparison, the PID response is also shown in red. The inset shows the normalized PID and deconvolved response to better compare the waveform of the two sensors. In order to test whether a major shift occurred due to the deconvolution procedure, we compared the threshold times (time to 5% of the max from the valve opening) of the deconvolved signal and the PID responses (**Fig. 1E**) for individual ethanol presentations. A linear relation (r = 0.9391, p<0.001) exists between the threshold time of the deconvolved signal and the PID responses.

**Figure 1.**
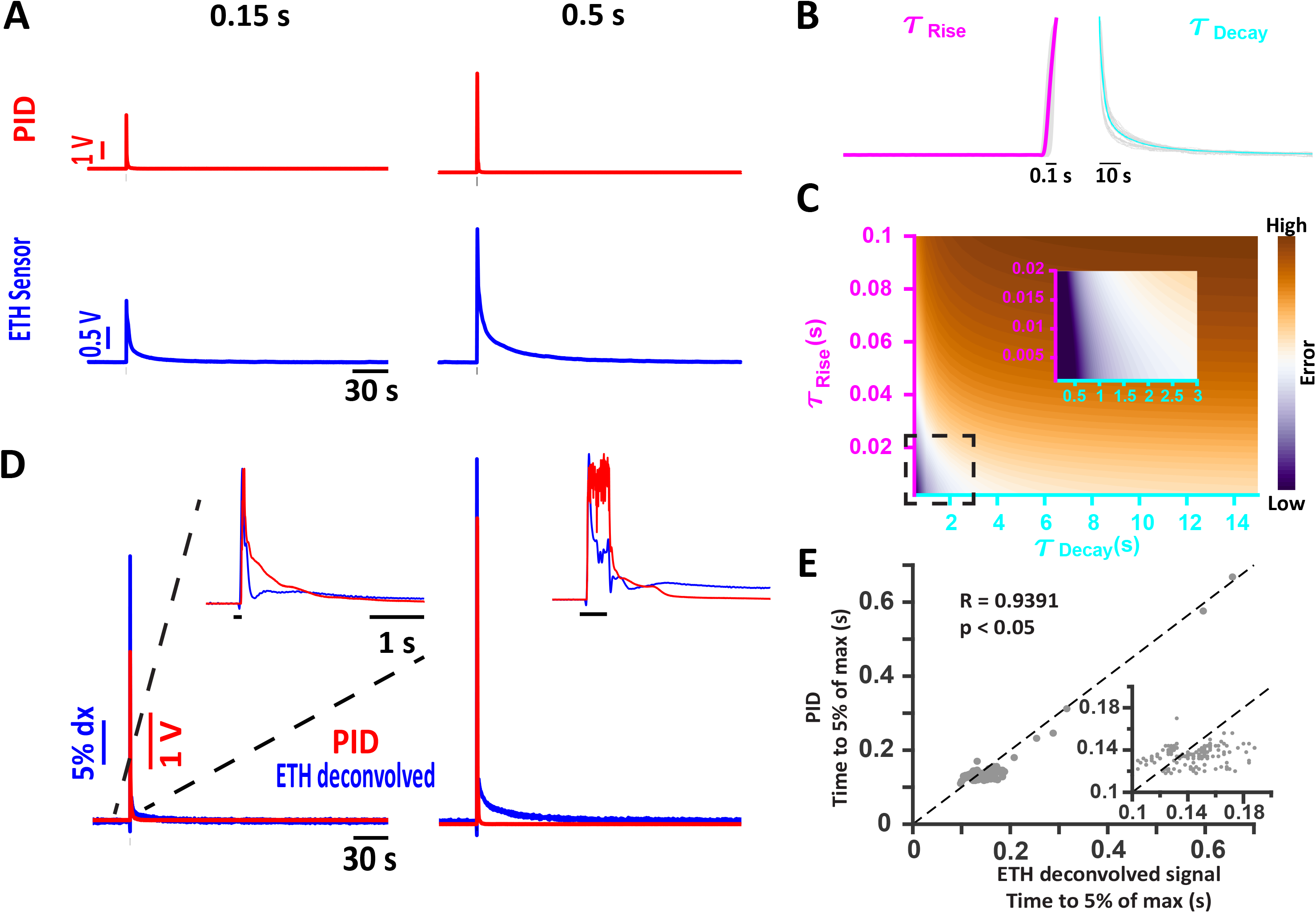
Ethanol sensor response can be deconvolved using a difference of two exponential kernel. **A)** Representative traces of the PID (red) and the ethanol sensor (blue) response to brief ethanol presentations. The duration of the ethanol pulse is indicated at the top of each column. **B)** Ethanol sensor responses from individual presentations (gray) were averaged to get an estimate of the rise time (τ_Rise_) to be around 0.06 s (magenta) and the decay time (τ_Decay_) to be around 3.5 s (cyan). **C)** Optimization of the deconvolution kernel by minimizing the mean error between the deconvolved ethanol responses and the PID responses (see Materials and Methods) using a family of kernels with varying τ_Rise_ and τ_Decay_ values. For later figures, the τ_Rise_ of the kernel was set at 0.002 s and the τ_Decay_ was set at 0.5 s. **D)** The deconvolved ethanol signal (blue) from the raw recordings of the ethanol sensor shown in **A** compared with the PID recordings. The inset shows the zoomed-in view of the normalized deconvolved ethanol response and the PID response to compare the waveforms of the PID response and the deconvolved ethanol signal. The black bar in the inset shows the duration of ethanol presentation. **E)** A linear relation between the threshold time (time from the valve opening to 5% of the max) between the deconvolved ethanol signals and the PID responses exists. Inset shows the zoomed-in view of the cluster of points in the ranges shown.

The summary data from all the durations equal to 0.3 s and less are presented in **Fig. 2A** as a heatmap where each row represents a single trial aligned with respect to the valve opening. The PID responses are presented in red, while the deconvolved ethanol sensor signals are presented in blue. Overlaying the two responses we can see coincident activity in magenta (right). Furthermore, the peak times (time of the maximum amplitude response with respect to the valve opening) of the deconvolved signal and the PID response (**Fig. 2B**) show a linear relationship (r = 0.8903, p < 0.001) indicating the coincident activity of the two sensors.

**Figure 2.**
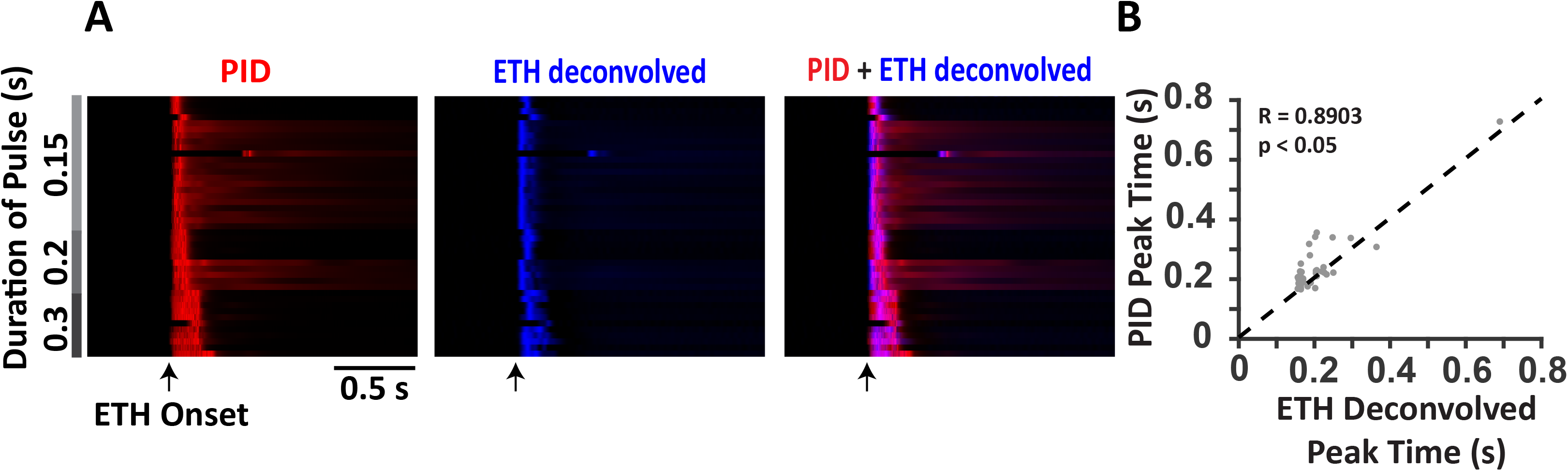
Peak times of the deconvolved ethanol signals coincide with the peak times of the PID responses for brief pulses of ethanol. **A)** Summary data of PID responses (red; left) and the deconvolved ethanol signals (blue; center) to brief durations of ethanol pulses (shown in gray) presented as a heatmap where each row is a single trial aligned with respect to the valve opening (arrow). Overlaying the deconvolved ethanol signals over the PID responses (right) shows coincidence of the peaks from the two signals (magenta). **B)** Linear relation between the peak times of the PID responses and the peak times of the deconvolved ethanol signals.

### Frequency characteristics of the ethanol sensor

We then tested the ranges of frequency at which the ethanol sensor can detect signals by conducting paired PID and alcohol sensor recordings to ethanol pulse stimuli of 5, 10 and 15 Hz frequencies. **Fig. 3A** shows the raw ethanol sensor and PID recordings for the frequencies tested. The raw sensor recordings were then deconvolved using the same kernel as used for the single pulses (see above). The deconvolved signal is shown in **Fig. 3B**. To better compare the deconvolved signal and the PID recordings, 1 s zoomed-in views of the signals are presented in **Fig. 3C** showing identical responses to the ethanol fluctuations.

**Figure 3.**
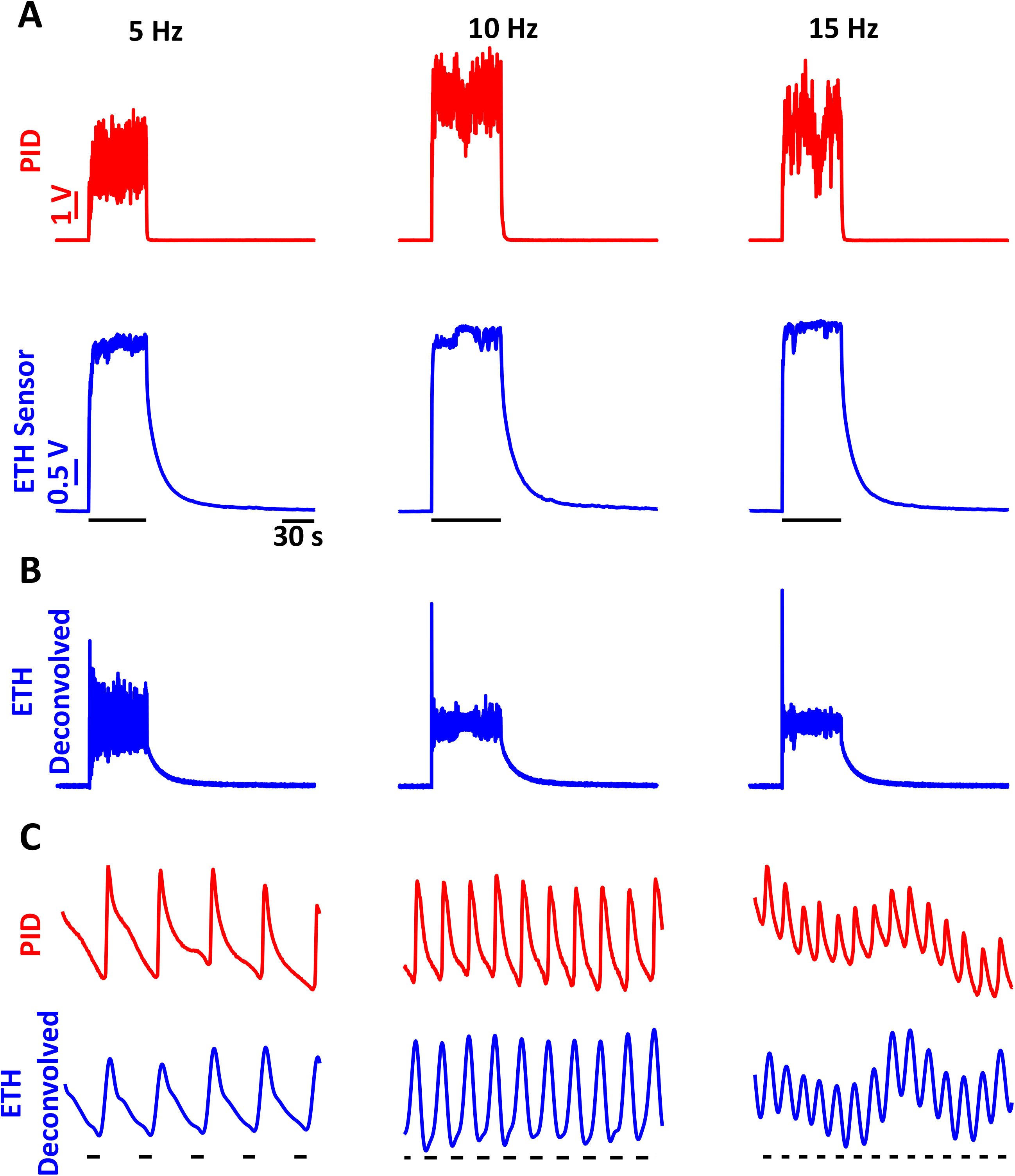
Frequency responses of the ethanol sensor show that the sensor can pick up frequencies of ethanol fluctuations up to 15Hz. **A)** PID (red) and ethanol sensor (blue) responses to ethanol fluctuations of 5 Hz (left), 10 Hz (center) and 15 Hz (right) frequencies. **B)** Deconvolved signals of the raw ethanol sensor recordings presented in **A**. **C)** A 1s zoomed-in view of the PID responses (red) and deconvolved ethanol signal (blue) showing near identical responses in the deconvolved signals and the PID recordings.

To better compare the two signals, we carried out a cross-correlation of the stimulation window and baseline windows as shown in **Fig. 4**. The mean cross-correlation (n = 5 for each frequency) shows peaks near the cycle times (dashed lines). There is a consistent small difference in the peak times of the mean cross-correlation with respect to the cycle times (Mean: 3ms) across all the frequencies which might be explained by the delay in the opening of the valves (Clippard EV-2-12, response time: 5-10ms).

**Figure 4.**
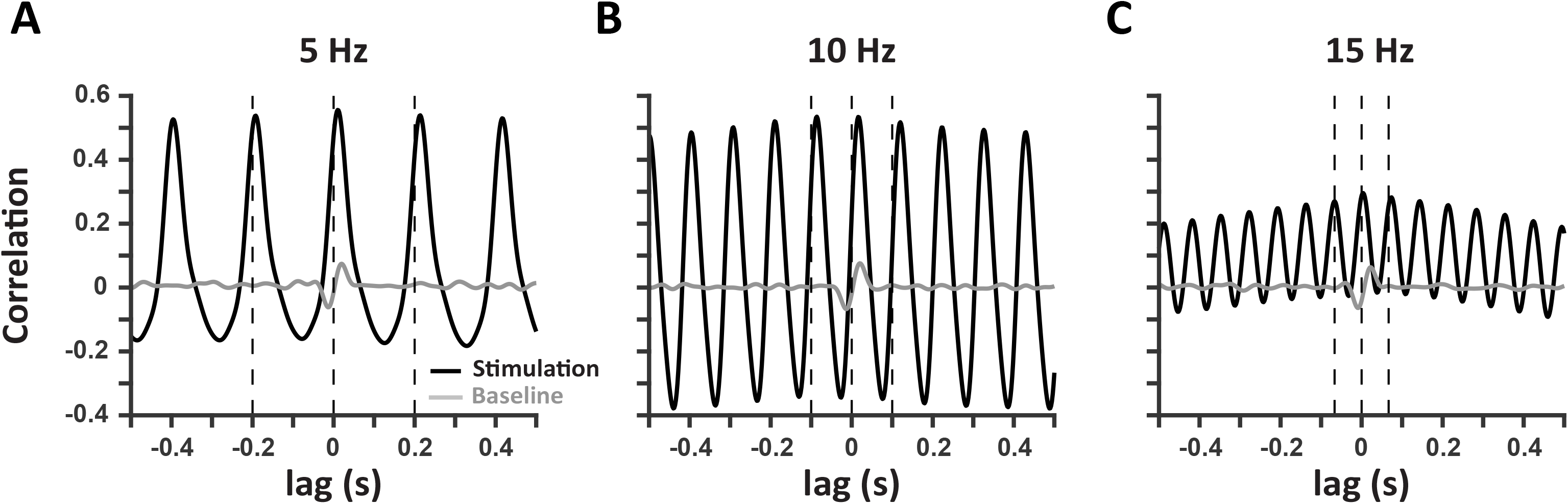
Deconvolved ethanol signals are correlated with the PID signals at multiple frequencies. Mean cross-correlation of the deconvolved ethanol signal with the PID signal during stimulation (black) at 5 Hz **(A**), 10 Hz (**B**) and 15 Hz (**C**) frequencies shows robust correlation as compared to during the baseline (gray). The dashed lines represent the cycle time for each frequency.

### Spatial characteristics of the ethanol sensor

To test how distance from the ethanol source affected the alcohol sensor reading and responses to a more behaviorally relevant odor stimulus, we next conducted paired PID and alcohol sensor recordings in a custom-made arena designed to create dynamic odor plumes (**Fig. 5A**). The recordings were carried out near the source (1), middle of the arena (2), and farther downwind (3) locations indicated. Representative traces from the three locations are shown in **Fig. 5B**. As can be seen, the increasing distance results in a decrease of the amplitude of both the PID and sensor readings. The mean correlations (Mean±SD: 0.460±0.175 (1); 0.4708±0.1584 (2); 0.4343±0.0937 (3)) between the PID and the deconvolved alcohol signal for the ethanol duration window are presented in **Fig. 5C**.

**Figure 5.**
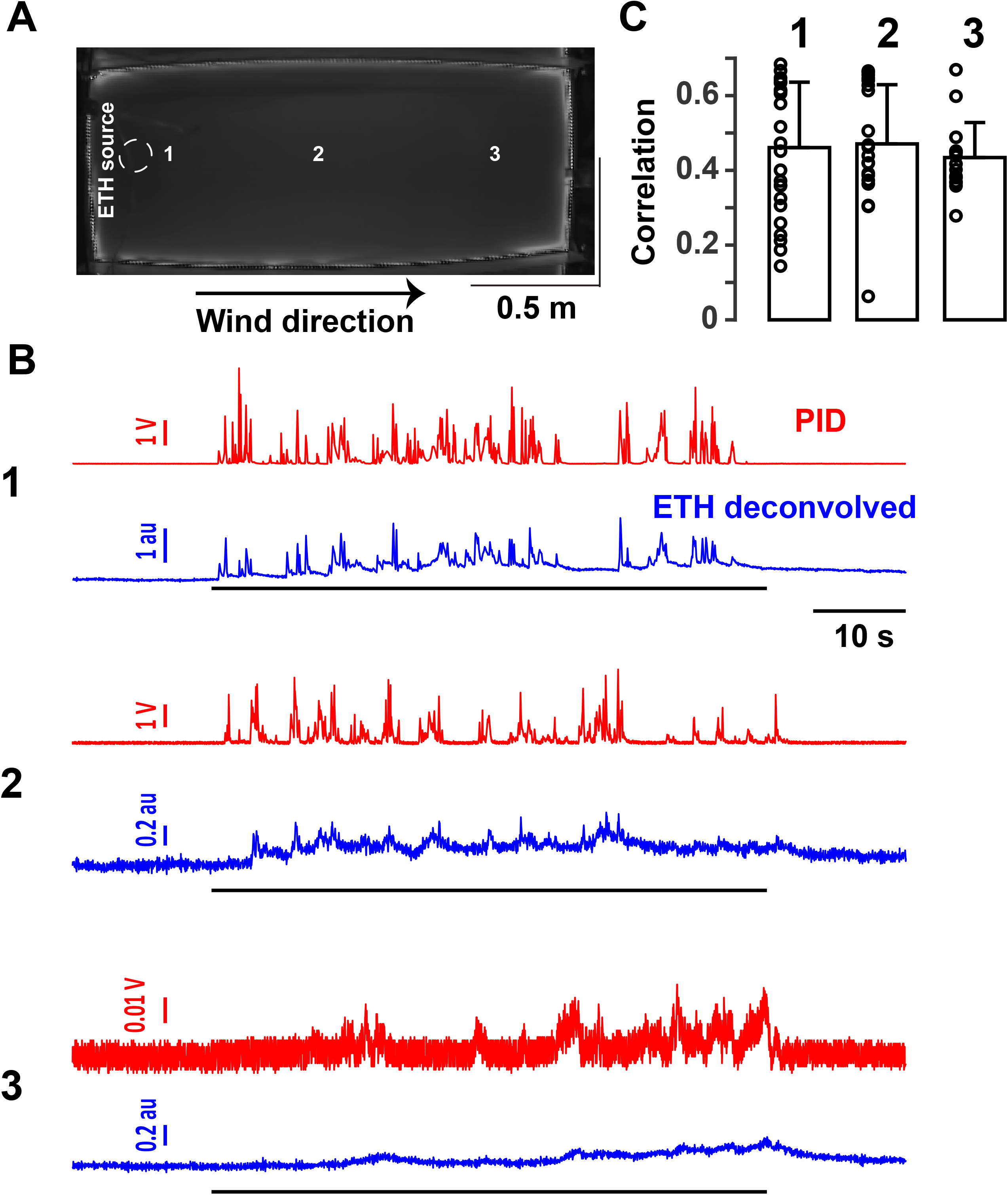
Deconvolved ethanol signals are correlated with the PID responses in turbulent airflow. **A)** An overhead image of a custom-designed arena (0.9*2*0.9 x*y*z m^3^) to create dynamic odor plumes. The dashed circle represents the location of the source of ethanol port while the arrow points out the wind direction. Also indicated are the approximate locations near the port (1), middle of the arena (2) and farthest downwind (3) where paired PID and ethanol sensor recordings were carried out. **B)** Representative traces of the PID (red) and deconvolved ethanol (blue) signals at the locations indicated in **A**. The black bars represent the ethanol stimuli. Notice the changing scales for the amplitude at farther distances from the ethanol port. **C)** Mean+SD correlations (bars) at locations near the port (1), middle of the arena (2), and farthest downwind (3) from the ethanol port. Circles are the correlations of individual trials.

### Behavioral and neurophysiological responses correlated with ethanol contacts detected by the alcohol sensor

To test if neural processing in the early olfactory pathways were correlated with sensor recordings, we conducted paired widefield calcium imaging over the dorsal surface of the olfactory bulb (OB) of head-fixed Thy1GCaMP6f (GP 5.11) mice [6] with ethanol sensor readings during ethanol plume presentations in a specially designed wind tunnel to create dynamic odor plumes. **Fig. 6A** shows the baseline image of the dorsal surface of the OB (left) with the standard deviation image (right) after ethanol plume presentation. Ethanol plume presentation resulted in glomerular activity patterns, obtained by cumulative non-negative matrix factorization (CNMF) [8], that were correlated with the deconvolved ethanol signal from the sensor. **Fig. 6B** shows the difference in fluorescence traces from 10 different glomeruli aligned with respect to the first peak of the deconvolved signal. As can be seen, the glomerular activity showed varied responses with some glomeruli following the fluctuations in the ethanol plume. **Fig. 6C** shows the mean activity of the glomeruli shown in **Fig. 6B** to 40 different presentations of the ethanol plume. The mean first peak of the ethanol signal detected by the sensor is also presented. As can be seen, the glomerular activity is consistent with the detection by the sensor. In order to better analyze how the population-level activity in the OB changed with the plume presentation, we conducted a PCA on the population-level activity in the OB. **Fig. 6D** shows the trajectory of the activity in the first two PC (accounting for 48% of the variance) space after the plume detection. As can be seen, the trajectory after the plume detection (blue line with red dots) moves away from the resting state (blue lines alone). The PCA was then carried out on the glomerular dynamics data from individual sessions and the Euclidean distance between all the PCs during the resting and post-plume detection was calculated. **Fig. 6E** shows that the distance between the PCs post-plume detection increases during 5 individual recording sessions. These results confirm that the plume detection by the alcohol sensor is correlated with neural processing in the OB.

**Figure 6.**
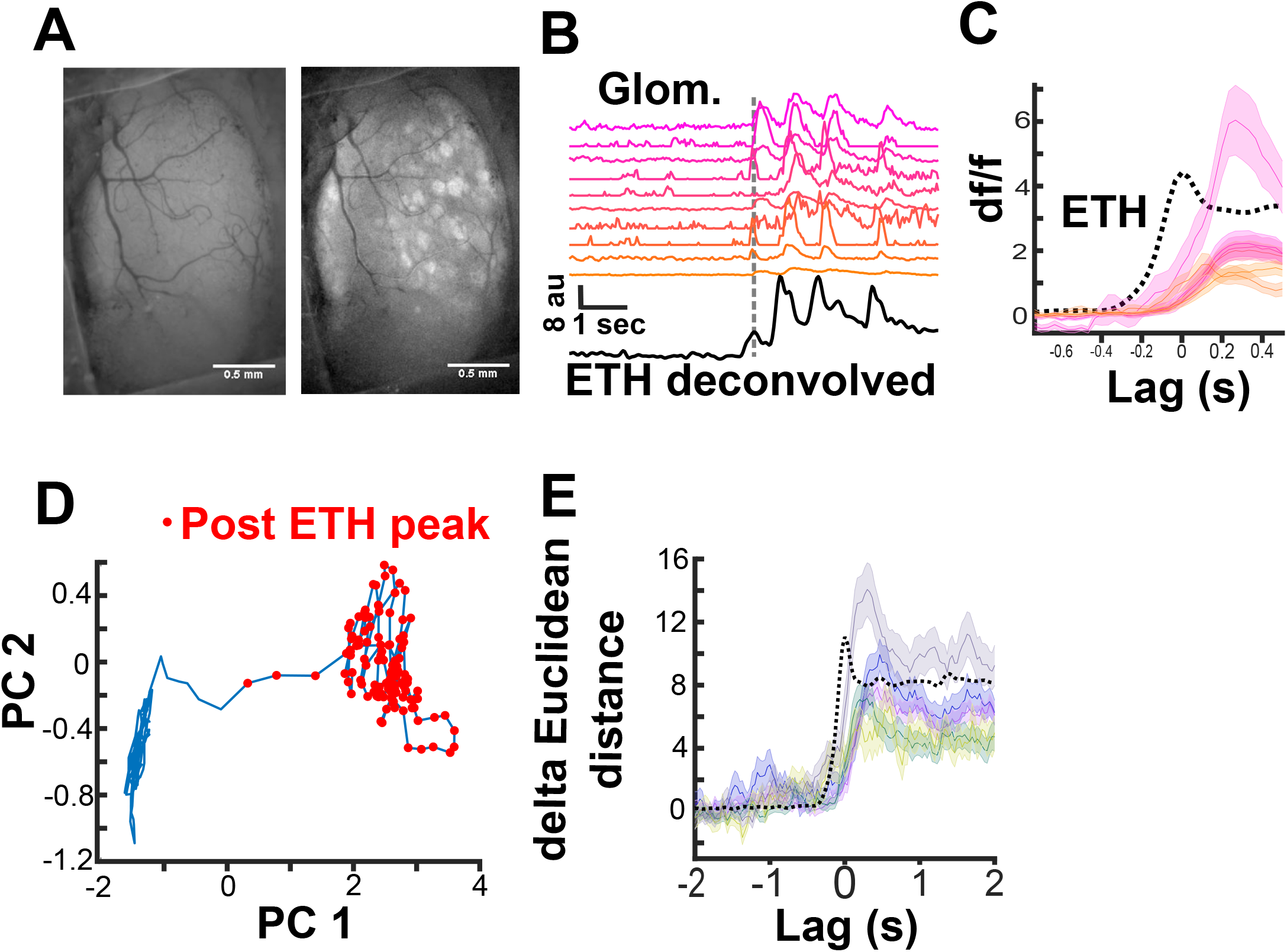
Ethanol plume detected by the ethanol sensor correlates with neural processing in the olfactory bulb. **A)** A representative cranial window over the dorsal surface of the olfactory bulb of a Thy1-GCaMP6f GP5.11 mouse. Left is at rest while right is the standard deviation image after ethanol plume exposure. **B)** Calcium traces from 10 different glomeruli aligned to the first peak of the ethanol plume (dashed line). **C)** Mean +/− SEM traces from the 10 glomeruli averaged over 40 different ethanol plume exposures zoomed in to focus the moment of plume contact. **D)** Movement in the first 2 PC (accounting 48% of the variance) space post ethanol plume exposure from the data presented in **B** and **C**. **E)** Sum of the Euclidean distance from rest across all the PCs aligned to plume detection from 5 different imaging sessions. Each trace is the mean +/− SEM across 40 ethanol plume presentations during each session.

To study how plume detection shapes behavior in plume-tracking mice, we trained water-deprived mice to associate ethanol with water. Next they were tasked to find the ethanol source in a large wind tunnel (0.9*2*0.9 x*y*z m^3^). The mice were equipped with head-mounted sensors and their behavior and ethanol plume contact were recorded. Two instances of a mouse engaged in this task are shown slowed about 3x the real time (see supplemental videos). In the video, the red marks the animal while the trail represents the positions of the animal in the last half second and the plus sign denotes the location of the ethanol source in the arena. The white trace is a moving median of the speed of the mouse, while the blue trace is the deconvolved ethanol signal. Plume detection (defined as points where the deconvolved signal exceeds 2 SD from the baseline marked as points where the red marker turns black) is associated with an increase in running speed and orientation toward the plume source.

## Discussion

Correlating real-time olfactory information with behavior and physiological recordings has been a challenging avenue due to the dynamic nature of the olfactory stimulus. Vickers and Baker (1994) first conducted EAG recordings from antennae mounted on freely behaving moths to correlate odor stimuli with behavior [9]. While EAG is an effective measure to monitor odor information, EAG suffers from degradation of signals over time. Another technology routinely used is the PID [5]. However, the PID sensor is bulky and not feasible for mobile monitoring. We therefore propose using metal oxide gas sensors, specifically alcohol sensors, to monitor the environment during behavioral and physiological recordings. These sensors are inexpensive and can be easily acquired.

One limitation of using metal oxide gas sensors is their long decay time to transient activation. We, therefore, designed kernels resulting from the difference of two exponentials to deconvolve the signal. The deconvolved signals showed a good correlation with the PID responses over time, frequency and spatial scales. We also show good correlation between neural responses in the OB of mice and the deconvolved signals from the sensor. In addition, behavioral changes from freely behaving animals in response to odor contacts detected by the sensor are presented. We, therefore, propose this method to be a robust and cost-effective method to study the odor-dependent changes in neural processing and behaviors in rodents, and can thus be of great value to olfactory neuroscientists.

